# Structure-function relation of the developing calyx of Held synapse *in vivo*

**DOI:** 10.1101/2020.01.07.893685

**Authors:** Martijn C. Sierksma, Johan A. Slotman, Adriaan B. Houtsmuller, J. Gerard G. Borst

**Author notes:** An earlier version of this article was first published as a preprint: Sierksma MC, Slotman JA, Houtsmuller AB, Borst JGG (2020). Structure-function relation of the developing calyx of Held synapse *in vivo*. bioRxiv. https://doi.org/10.1101/2020.01.07.893685.

## Abstract

In adult rodents, a principal neuron in the medial nucleus of the trapezoid (MNTB) is generally contacted by a single, giant axosomatic terminal called the calyx of Held. How this one-on-one relation is established is still unknown, but anatomical evidence suggests that during development principal neurons are innervated by multiple calyces, which may indicate calyceal competition. However, *in vivo* electrophysiological recordings from principal neurons indicated that only a single strong synaptic connection forms per cell. To test whether a mismatch exists between synaptic strength and terminal size, we compared the strength of synaptic inputs with the morphology of the synaptic terminals. *In vivo* whole-cell recordings of the MNTB neurons from newborn Wistar rats of either sex were made while stimulating their afferent axons, allowing us to identify multiple inputs. The strength of the strongest input increased to calyceal levels in a few days across cells, while the strength of the second strongest input was stable. The recorded cells were subsequently immunolabeled for vesicular glutamate transporters (VGluT) to reveal axosomatic terminals with structured-illumination microscopy. Synaptic strength of the strongest input was correlated with the contact area of the largest VGluT cluster at the soma (*r* = 0.8), and no indication of a mismatch between structure and strength was observed. Together, our data agree with a developmental scheme in which one input strengthens and becomes the calyx of Held, but not with multi-calyceal competition.

**Key points summary:**

1. During development the giant, auditory calyx of Held forms a one-to-one connection with a principal neuron of the medial nucleus of the trapezoid body.
2. While anatomical studies described that most of the target cells are temporarily contacted by multiple calyces, multi-calyceal innervation was only sporadically observed in *in vivo* recordings, suggesting a structure-function discrepancy.
3. We correlated synaptic strength of inputs, identified in *in vivo* recordings, with post hoc labeling of the recorded neuron and synaptic terminals containing vesicular glutamate transporters (VGluT).
4. During development only one input increased to the level of the calyx of Held synapse, and its strength correlated with the large VGluT cluster contacting the postsynaptic soma.
5. As neither competing strong inputs nor multiple large VGluT clusters on a single cell were observed, our findings did not indicate a structure-function discrepancy.

## Introduction

Synapses are specialized structures that allow presynaptic signals to be transmitted to the postsynaptic side. Most central synapses use neurotransmitters for signaling, and these chemical synapses come in a large variety of sizes and shapes. While most synapses have a diameter of about one μm and are formed on a dendrite, some synapses can be >10 μm, and encompass >100 release sites (Walmsley *et al*., 1998; Atwood & Karunanithi, 2002; Rollenhagen & Lübke, 2006). These large synapses have facilitated the study of the biophysical properties of synapses. A prime example is the calyx of Held synapse in the auditory brainstem. The presynaptic calyx spans ^~^20 μm, large enough to be accessible for patch-clamp electrophysiology (Forsythe, 1994; Borst *et al*., 1995) and to allow simultaneous recording of the calyx and its postsynaptic target (Borst *et al*., 1995), a glycinergic neuron in the medial nucleus of the trapezoid body (MNTB). Owing to the presence of hundreds of active zones (Sätzler *et al*., 2002; Taschenberger *et al*., 2002; Dondzillo *et al*., 2010), the calyx of Held can rapidly trigger postsynaptic action potentials (APs), thus functioning as a fast, high-fidelity, inverting relay in the auditory brainstem (Borst & Soria van Hoeve, 2012).

During its development, the calyx of Held synapse grows from an axon of a globular bushy cell (GBC) that initially forms small boutons. In rodents these contacts already appear before birth (Kandler & Friauf, 1993; Hoffpauir *et al*., 2006; Rodríguez-Contreras *et al*., 2008; Hoffpauir *et al*., 2010; Holcomb *et al*., 2013). Although the innervation initially is highly divergent, eventually a GBC will give rise to only one or a few calyces, and most adult principal neurons are innervated by a single calyx of Held (Held, 1893; Morest, 1968; Kuwabara *et al*., 1991; Kandler & Friauf, 1993; Rodríguez-Contreras *et al*., 2006). The developmental mechanisms that ensure the transition to the one-on-one innervation are largely unclear (Yu & Goodrich, 2014). This transition happens between the second and the fifth postnatal day (P2-5). At P5 the strength of one input overshadows the other ones (Chuhma & Ohmori, 1998; Hoffpauir *et al*., 2006; Hoffpauir *et al*., 2010; Sierksma *et al*., 2016), and one of the terminals has expanded over the soma of the neuron (Hoffpauir *et al*., 2006; Holcomb *et al*., 2013). From these studies it was suggested that half of the MNTB neurons are contacted by multiple calyces at this age range (Hoffpauir *et al*., 2010; Holcomb *et al*., 2013; Milinkeviciute *et al*., 2019), in agreement with observations from mouse slice recordings (Bergsman *et al*., 2004; Hoffpauir *et al*., 2010; Xiao *et al*., 2013). Note that multi-calyceal innervation is defined here as an MNTB neuron that is innervated by more than one calyx of Held, and not as an MNTB neuron that is innervated by multiple calycigenic axons.

The developmental changes that underlie the formation of the calyx of Held synapse may involve activity-dependent processes. In newborn, pre-hearing rodents, spontaneous activity, arising in the cochlea (Tritsch *et al*., 2010), propagates across the developing auditory brainstem (Sonntag *et al*., 2009; Tritsch *et al*., 2010; Crins *et al*., 2011; Sierksma *et al*., 2016). The correlated, bursting activity of presynaptic terminals (Sierksma & Borst, 2017) causes plateau-like depolarizations in the postsynaptic cell (Sierksma *et al*., 2016) and bursting activity (Tritsch *et al*., 2010; Crins *et al*., 2011; Clause *et al*., 2014) which may induce synaptic plasticity. The presence of multiple calyces contacting the same target neuron would suggest that the formation of a one-to-one connection involves some form of competition during which one calyx rapidly expands and increases in strength, whereas the other contacts retract and decrease in strength. However, in the rat, *in vivo* physiological evidence for calyceal competition was observed in only very few cases (Sierksma *et al*., 2016).

These findings may indicate a discrepancy between terminal size and synaptic strength. To test this possibility, one needs to correlate synaptic structures with synaptic strength. Serial sectioning of tissue combined with EM offers the possibility of fully reconstructing every synapse in a restricted volume (Denk & Horstmann, 2004; Hoffpauir *et al*., 2006; Holcomb *et al*., 2013), and allows the reconstruction of entire calyces (Sätzler *et al*., 2002; Holcomb *et al*., 2013; Qiu *et al*., 2015). However, it does not offer a direct estimate of synaptic strength. Whereas slice electrophysiology has the advantage of allowing simultaneous recordings of pre- and postsynaptic structures (Borst *et al*., 1995; Rodríguez-Contreras *et al*., 2008), there is the uncertainty associated with possible cutting of inputs during slice preparation. Immunolabeling of vesicular glutamate transporters (VGluT) localizes synapses in the MNTB (Rodríguez-Contreras *et al*., 2006; Rodríguez-Contreras *et al*., 2008; Soria Van Hoeve & Borst, 2010; Milinkeviciute *et al*., 2019). Parvalbumin may identify calyces, but not at the youngest ages (Lohmann & Friauf, 1996, but see Felmy & Schneggenburger, 2004), and its presence does not indicate whether the calyx is release-competent. As Piccolo, an active zone protein (Südhof, 2012; Gundelfinger *et al*., 2016), is present in young calyces (Dondzillo *et al*., 2010), the combination of VGluT and Piccolo may indicate synaptic connectivity in the MNTB. How VGluT relates to synaptic strength is still unclear.

Here, we combined VGluT immunolabeling with *in vivo* whole-cell recordings during which we recorded the inputs that are regularly active (Lorteije *et al*., 2009; Sierksma *et al*., 2016). We applied electrical stimulation of the afferent axons to further identify the developing inputs. The recorded cells were subsequently immunolabeled for VGluT and Piccolo, allowing a direct comparison between synapse strength and structure for the developing calyx of Held synapse.

## Methods

### Ethical approval

All procedures conformed to the European Directive 2010/63/EU and were approved by the local animal ethics committee (EDC, project no. 115-14-11). Timed-pregnant Wistar rats (WU) were purchased from Charles River and housed within the Erasmus animal facility (EDC). The day of birth or finding the litter was taken as postnatal day (P)0. The dams had *ad libitum* access to food and water, and additional bedding and shelter material was provided for nest building. Wistar pups of undetermined sex were taken from the nest on the day of the experiment. The authors understand the ethical principles under which the journal operates and confirm that their work complies with the animal ethics checklist provided by the editorial board.

### Surgery

The pup was anesthetized with isoflurane administered together with medical oxygen. Anesthesia depth was checked by the paw pinch prior to the surgery. We used a ventral approach to gain access to the auditory brainstem, as described previously (Rodríguez-Contreras *et al*., 2008; Sierksma *et al*., 2016). After recording the animal was deeply anesthetized and perfused with 6-10 mL of cold saline (0.9% NaCl, w/v), followed by 8-10 mL of 4% of paraformaldehyde (PFA, w/v) dissolved in 0.12 M phosphate-buffer (PB, pH 7.2-7.4).

### In vivo electrophysiology

First, we searched for bursting activity with a glass electrode to find the presumed location of the MNTB. When bursting activity was found, the search pipette was retracted and a bipolar stimulation electrode (MicroProbes for Life Science, PI2ST30.1H10) was placed contralateral from the recorded MNTB, straddling the trapezoid body, as described by Crins *et al*. (2011). Successful placement was assessed by measuring the stimulation current threshold and the presence of a field potential.

Whole-cell recordings with biocytin (2 mg/mL) in the pipette solution were made from principal cells; to be confident that the recording was obtained from the same neuron as the one that was recovered with immunolabeling, we made whole-cell recordings from a single principal neuron per animal. Whenever a glia cell or non-principal cell was encountered, the pipette was retracted and a new pipette was used. Membrane potentials were compensated prior to the recordings for a liquid junction potential of −11 mV. Drift in membrane potential was <5 mV and remained uncorrected. Recordings were made with a MultiClamp 700B in current-clamp mode, with bridge balance set in the range of 20-70 MΩ and pipette capacitance compensation ^~^4-6 pF. Signals were low-pass filtered with a 4-pole Bessel filter at 10 kHz and digitized by a DigiData 1440A (Molecular Devices Co.) at 25 kHz. Acquisition was done with Clampex 10.2 running on a Windows XP computer.

Stimulation intensities were manually adjusted during the experiments. First, we quickly assessed at which stimulation levels new excitatory postsynaptic potentials (EPSPs) were recruited, and at which polarity most inputs could be discerned. Then, stimulation intensity was lowered to the activation threshold of the first input, and step-wise increased over time. In some experiments the stimulation intensity was also stepwise reduced, and then the data were pooled. For every stimulation strength >30 sweeps were collected at 2 Hz. Stimulation current was typically below 0.4 mA to prevent damage to the axons; at high stimulation currents (0.3 mA or higher) antidromic APs with latencies <1 ms were often elicited in the principal neuron. This could still happen even though we used bipolar stimulation electrodes, placed on the contralateral side from the recorded neuron. Preferably, the stimulation electrode would be placed at the midline, where all calycigenic axons converge ventrally. However, to avoid direct postsynaptic activation we placed the stimulation electrode more laterally. This location may have led to preferential activation of the axon bundles that run ventrally from the other MNTB (Kandler & Friauf, 1993).

Labeling calyces by *in vivo* whole-calyx recordings or electroporation was performed as described previously (Sierksma & Borst, 2017).

### Antibodies

The following primary antibodies were used: rabbit polyclonal anti-Piccolo (Synaptic Systems 142003, RRID: AB_2160182; 1:1000), guinea-pig polyclonal anti-vesicular glutamate transporter 1 (Millipore AB5905, RRID: AB_2301751; 1:3000) and 2 (Millipore AB2251, RRID: AB_1587626; 1:3000). Secondary antibodies Alexa Fluor 488 against rabbit and Alexa Fluor 645 against guinea pig were obtained from Jackson (1:200 or 1:400). Streptavidin-Alexa Fluor 594 conjugate (1:200) was obtained from Thermo Fisher Scientific. Throughout, VGluT labeling refers to both VGluT1 and VGluT2.

### Immunolabeling procedure

Immunolabeling procedure was based on the free-floating method. After perfusion the brain was carefully removed from the skull, and post-fixed overnight at 4 °C. Then, it was left overnight in 10% (w/v) sucrose in 0.1 M PB, embedded in 10% (w/v) gelatin and 10% (w/v) sucrose, and again fixed overnight at 4 °C in 30% (w/v) sucrose and 10% (w/v) formaldehyde. The brain was cryoprotected with 30% (w/v) sucrose solution in 0.1 M PB for >24 hr at 4 °C. Coronal slices (40 μm) were made on a freezing microtome, and were collected in 0.1 M PB. The sections were heated for 3 hr to 80 °C in 10 mM sodium citrate (pH 6 at RT), washed four times for 10 min with 0.9% (w/v) NaCl in 0.05 M PB (PBS). Then, the sections were pre-absorbed for 1 hr at RT with 10% (v/v) normal horse serum and 10% (v/v) Triton X-100 in PBS followed by 36-48 hr incubation under gentle agitation at 4 °C with primary antibody solution containing 2% (v/v) normal horse serum, 0.4% (v/v) Triton X-100 and the primary antibodies. The slices were washed four times for 10 min in PBS at RT, and incubated for 2 hr under gentle agitation at RT in the secondary antibody solution containing 2% (v/v) normal horse serum, 0.4% (v/v) Triton X-100, and the secondary antibodies. For the biocytin-filled cells the streptavidin conjugate was also added at this step. The slices were washed once for 10 min at RT with PBS, and incubated for 10 min RT in 0.3 μM DAPI (D3571, Invitrogen) in 0.1 M PB. Sections were then washed three times at RT with 0.1 M PB, mounted on glass coverslips with gelatin-chrome alum, air-dried, and closed with Mowiol mounting solution containing 10% (w/v) Mowiol 4-88, 25% (v/v) glycerol in 0.1 M Tris-buffer (pH 8.5). Sections were kept at 4 °C in the dark until further use.

### Confocal and structured illumination imaging

Confocal imaging was done on a Zeiss LSM 700 microscope equipped with a plan-apochromat 20x, 0.75 NA and a 63x, 1.4 NA objective, and PMT detectors, or on a Zeiss Elyra PS1 microscope, which is described below. Confocal images were acquired with optimized settings for laser power, detector gains and pin hole diameters. Large (2048×2048 pixels), high-resolution images were acquired with a pixel size of 0.041 μm laterally and 0.110 μm radially to facilitate the comparison with structured-illumination microscopy (SIM) images. Low-resolution tile images were acquired with a pixel size of 0.274 μm.

Structured-illumination imaging was done on a Zeiss Elyra PS1 system equipped with 488, 561 and 642 nm, 100 mW diode lasers; fluorescence was acquired with a Zeiss plan-apochromat 63x, 1.4 NA objective and an Andor iXon DU 885 EMCCD camera (1002 x 1004 pixels). Gratings were presented at 5 phases and 5 rotations for every depth. Sampling interval was 110 nm axially and the reconstructed super-resolution image has a pixel size of 41 nm. The fluorophore signals were acquired sequentially. Reconstructions were made with built-in algorithms using ZEN 2012 SP1 black (v8.1.3.484).

### Electrophysiological analysis

EPSPs were detected as described previously (Sierksma *et al*., 2016) using an EPSP minimal rate of rise (>0.7 V/s) and an EPSP onset based on the third derivative. EPSPs that were present between 1 to 8 ms post-stimulation were categorized as ‘evoked’ EPSPs. We grouped the evoked EPSPs based on stimulus strength, latency and rate of rise. We assumed that with increasing stimulation intensities previously-recruited inputs would still be activated. For consecutive stimulus intensities Welch’s *t* was calculated for the EPSP latency and the rate of rise, and both *t* values were added together. When the summed *t* value was above 6 and the increase of the response rate of rise was >20%, we considered the evoked EPSPs to differ from each other. We visually checked whether this difference was due to a new input or just a shift in activation probability of a previously activated input. For a new input its EPSP rate of rise was calculated by subtracting the EPSP rates of rise of previously identified inputs when they were of similar latencies (±0.5 ms). For four recordings we found EPSPs with longer latencies (Figure 1C), and for these recordings the Welch’s *t* test was calculated but restricted to a latency domain defined by the experimenter. For the example in Figure 1C this was 1-3 ms. Lastly, the EPSP rate of rise of an input was checked against the distribution of spontaneously-occurring EPSP rates of rise, and the defined input was only accepted if spontaneous EPSPs with similar rates of rise were recorded.

**Figure 1.**
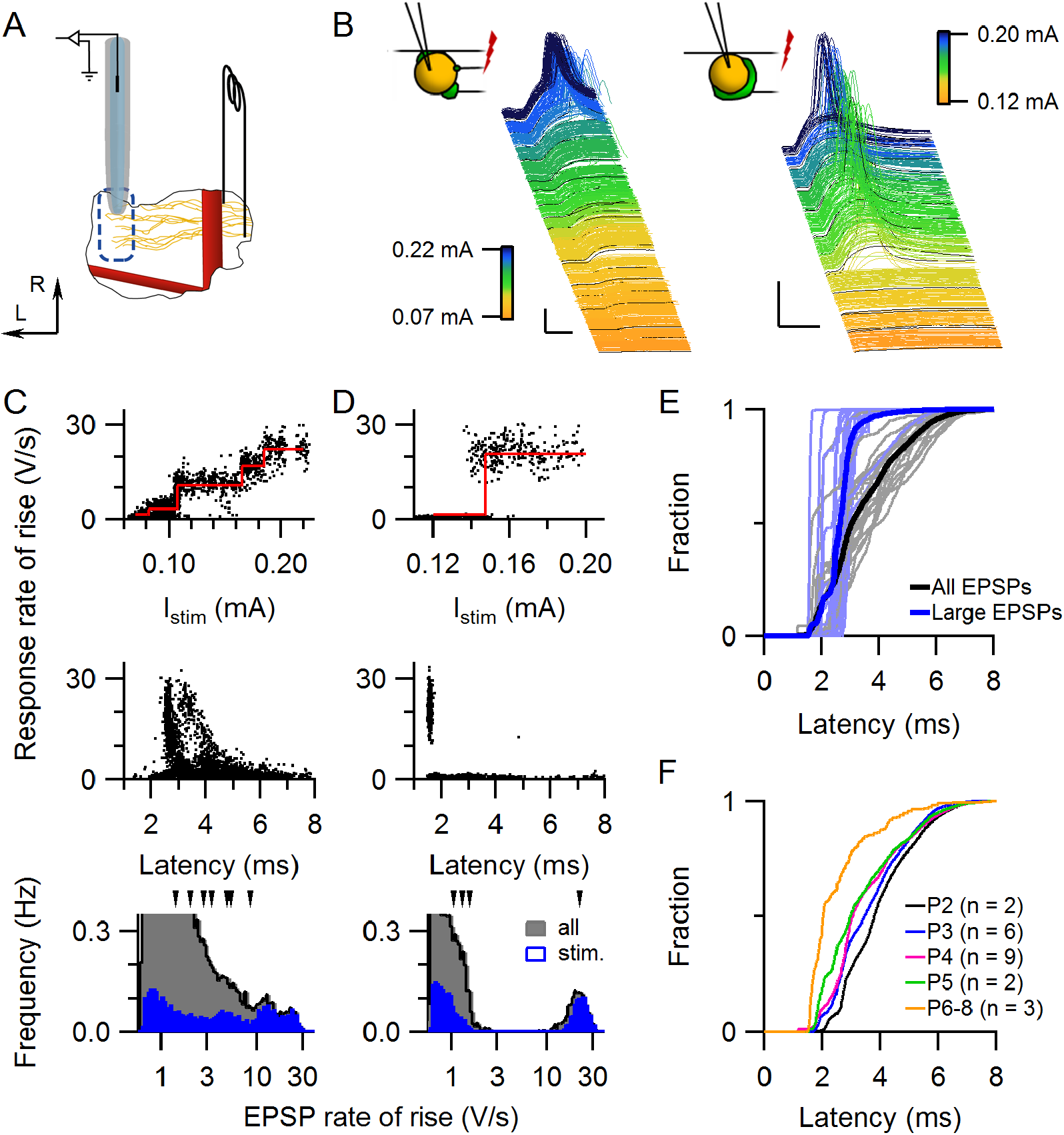
Responses to current stimulation of developing calycigenic axons. (A) Schematic drawing of the experimental approach. A bipolar stimulation electrode straddling the trapezoid body was positioned contralaterally from the recorded MNTB, and whole-cell recordings were made from principal cells of the MNTB. The basilar artery (rostrocaudal, thick red line) and the anterior-inferior cerebellar artery (mediolateral, red line) served as landmarks. Arrows indicate lateral (L) and rostral (R) directions. (B) Waterfall plots of two whole-cell recordings, sorted on stimulation intensity (color-coded as indicated in calibration bars). In the P4 example (left) a gradual increase in the evoked EPSP was observed, suggesting the recruitment of multiple inputs, while in the P6 example (right) a single, large EPSP was observed, which elicited a postsynaptic AP. Schematic drawing depicts the assumed synaptic innervation. Scale bars indicate 25 mV and 2 ms. (C) Identification of synaptic inputs. Rates of rise of evoked EPSPs vs. stimulation intensity (top) or EPSP latency (middle) for the P4 example shown in B. The bottom graph shows the histograms of the rates of rise of evoked and spontaneous EPSPs. Arrow heads on top of the graph indicate the rates of rise of identified inputs. Data points in the top graph were horizontally scattered by <50 μA for visualization purposes. (D) As C, except for the P6 example shown in B. (E) Cumulative distribution of latencies of all evoked EPSPs (black) and of large EPSPs (>10 V/s, blue). Individual cells are shown in light blue and grey. Large EPSPs have a shorter latency than most other EPSPs. (F) The average cumulative distribution of the evoked EPSP latencies grouped by age suggests a developmental increase in propagation speed.

In addition to the inputs identified by stimulation, we identified inputs from the spontaneous EPSPs. By comparing the distribution of EPSP rates of rise of all detected EPSPs in a recording with the evoked EPSPs, we could sometimes identify additional strong inputs (8 strong inputs in 8 cells). Within the spontaneous EPSPs, many had a small size (0.7-2 V/s), and these inputs were therefore hard to distinguish individually. If small EPSPs were not evoked, we visually selected peaks from the distribution of the EPSP rate of rise (Figure 1C-D) to define these smaller inputs, and added them to the dataset to acknowledge their existence. However, the inputs with small EPSPs were hard to demarcate and therefore likely underestimated.

Based on the EPSP size analysis, we identified on average 5.6 inputs per cell. For comparison, prior to synaptic elimination 5-12 axons contact a principal neuron in EM studies of the mouse MNTB (Hoffpauir *et al*., 2006). Another measure for how well we can identify all inputs can be obtained by comparing spontaneous and evoked inputs. In 8 out of 22 cells we were unable to activate the strongest input that was observed among the spontaneous activity, indicating that on the order of one in three inputs (8 out of 22 strongest inputs) were not electrically stimulated. Spontaneous activity partially negated this problem, but may have introduced a bias towards the most active and stronger inputs (see also below). To what extent synaptic inputs of the principal neuron are active in every spontaneous activity burst is unknown, but anecdotal evidence showed extensive co-occurrence of two distinct, large inputs during spontaneous bursts (Sierksma *et al*., 2016).

We tested whether the strong inputs could be further subdivided based on the presence of inter-event intervals that were shorter than the refractory period or the silent period within minibursts (Sierksma *et al*., 2016), but this did not lead to the identification of additional connections. Altogether, we believe that our combined analysis of spontaneous and evoked responses identified most synaptic inputs, except for some smaller inputs.

### Image analysis

Analysis of 3D-SIM image stacks was done in ImageJ using the FIJI framework (Schindelin *et al*., 2012; Schneider *et al*., 2012), making use of two custom-made 3D segmentation ImageJ plugins (available at: https://github.com/ErasmusOIC/FloodFill3D, https://github.com/ErasmusOIC/Sphere_Floodfill). In brief, we first segmented the cell body using the biocytin signal with the Sphere_Floodfill plugin. The biocytin signal was scanned with a 400 nm (10 pixels)-radius sphere. If 40% of the sphere contained intensity values above threshold, its center was assigned to the cell body. To segment the VGluT signal in 3D, we used the FloodFill3D plugin which scans all 26 neighboring pixels and assigns them to the VGluT structure if they are above an intensity threshold determined using the IsoData-method of ImageJ. To account for the width of the synaptic cleft, the cell body was enlarged by dilation. The amount of dilation (range 160-760 nm) was determined for each cell by maximizing the fraction of the dilated cell body covered by the VGluT structures. Where needed, axonal biocytin signals were manually removed from the cell body. The fraction of the surface area of the cell body that was covered by the VGluT clusters was calculated.

All images were pseudo-colored and contrasted in ImageJ. Color gamma enhancement was performed with Adobe Photoshop 19.1.0. We noticed that the peak intensities of Piccolo and VGluT did not seem to overlap in SIM images. From line profiles it became clear that Piccolo labeling did overlap with VGluT at lower intensities, but not at the highest VGluT intensities.

### Statistics

No blinding was applied to the analysis. Linear regression analyses against pup age were performed for the developmental effects. When the developmental effect was significant, the slope of the fit is reported; when not, the average is reported, and subsequently the *F*-statistic with the degrees of freedom, and the *p*-value are reported. Subsequent testing on the same data was Bonferroni-corrected, and are reported as ‘corrected *p*’. Correlations are reported as Pearson’s *r* or Spearman’s ρ after rank transformation. To compare prespike-associated strong inputs with the other strong inputs, we performed Welch’s *t*-test for the EPSP amplitude and latency. EPSP rate of rise and synaptic efficacy to trigger postsynaptic firing (P_AP_) were correlated with age. Age correction was performed by adding age as an explanatory variable to the regression analysis. P_AP_ was judged to have non-normally distributed residuals and therefore the regression analysis was done on the ranks. The *F*-statistic of the regression and the effect of prespike variable within this regression are reported. Sigmoid and linear fits were performed with Igor Pro 6.37 (WaveMetrics Inc.). To correlate the competition index with relative coverage we used Spearman’s ρ. *p*-values < 0.05 were considered significant.

## Results

### *Postsynaptic responses of* in vivo *stimulated fibers*

We made *in vivo* whole-cell recordings from principal neurons in the rat MNTB around the time the calyx of Held synapse forms. A bipolar stimulation electrode was placed contralaterally from the recording site to distinguish inputs based on their threshold for axonal action potential generation (see Methods; Figure 1A). Above a threshold of 83 ± 19 μA (mean ± SD, *n* = 30 animals, range: 65-140 μA) a field potential could be evoked in the MNTB. In a total of 22 MNTB cells, EPSPs could be evoked, and with increasing current stimulation we typically observed either the presence of a large, all-or-none, calyceal-like input, or more graded increases in EPSP size (Figure 1B). In both cases the evoked responses often could reach AP threshold (*n* = 16 of 22 cells). To identify the different inputs, we looked for discontinuities in the rates of rise and/or the latency of the evoked responses following an increase in stimulus intensity, as detailed in the Methods; these discontinuities were interpreted as the recruitment of a new input. A comparison of the rates of rise of the evoked EPSPs with those of the spontaneously-occurring EPSPs was used as a further check on the accuracy of the identified inputs (Figure 1C-D). In 8 out of the 22 cells we were unable to observe the largest spontaneous EPSPs following the electrical stimulation, suggesting that we failed to activate the strongest input for these cells. EPSP subpopulations were found with typical latencies of 1-3 and 4-8 ms (Figure 1E), as described in slice studies (Banks & Smith, 1992; Hoffpauir *et al*., 2006; Marrs & Spirou, 2012). The large EPSPs always belonged to the subpopulation with 1-3 ms delays (Figure 1E). On average the EPSP latencies became smaller during the first postnatal week (Figure 1F). In addition to early myelination (Rozeik & Von Keyserlingk, 1987; Hamano *et al*., 1996), an increase in axon diameter and/or ion channel densities may have contributed to this decrease in latency. The different subpopulations may reflect different developmental stages among the calycigenic axons.

Within this developmental period the principal neuron passes through a stage of exuberant synaptic connectivity (Holcomb *et al*., 2013). In addition to the 22 cells in which we recorded both evoked and spontaneous EPSPs, we recorded in 10 cells only the spontaneous activity. In total, we identified 180 putative inputs in 32 cells, 5.6 ± 1.6 per cell (mean ± SD). The distribution of input rate of rise was skewed to the right (Figure 2A). By sorting all inputs on their mean rate of rise, a discontinuity was observed at about 10 V/s (Figure 2A-B), which we used as a threshold to distinguish between weak and strong EPSPs. To have a measure of synaptic competition, we calculated the competition index, defined as the ratio of the rates of rise of the second strongest and the strongest input (Figure 2C). The competition index ranges from 1 to 0, where 1 indicates an equal rate of rise of the two strongest inputs, while if the competition index approaches 0 the strongest input has outcompeted the second strongest input in strength. The competition index decreased during development (−0.07 ± 0.02 day^−1^, n = 32 cells, *F*_1,30_ = 11, *p* = 0.002). This decrease can signify both an increase of the rate of rise of the strongest input and a reduction for the second strongest input. We therefore plotted the two against each other (Figure 2D). The rate of rise of the strongest input clearly increased during development (4.2 ± 1.1 V s^−1^ day^−1^, *F*_1,30_ = 14, corrected *p* = 0.001), while the rate of rise of the second strongest did not change (3.7 ± 0.2 V/s, *F*_1,30_ = 2.1, corrected *p* = 0.3). This was not related to a difference between evoked and spontaneous inputs, because inputs, whether evoked or identified within spontaneous activity, had similar rates of rise for both the strongest (stim.: 19 ± 10 V/s, n = 14 cells; spont.: 16 ± 12 V/s, n = 18 cells) and the second strongest input (stim.: 3.6 ± 1.2 V/s, n = 14; spont.: 3.7 ± 1.2 V/s, n = 18). In contrast to what may have been expected for competing inputs, the developmental increase in strength of the strongest input did not coincide with a decrease in the second strongest input.

**Figure 2.**
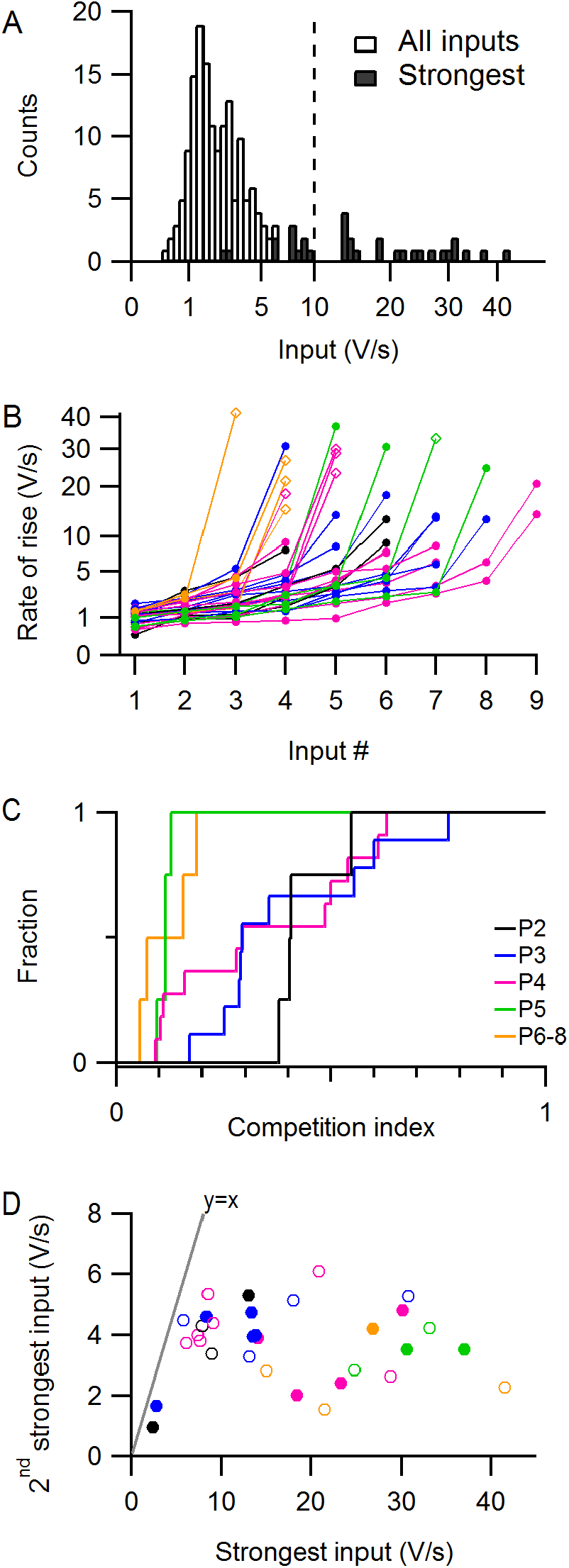
Synaptic competition in MNTB follows the expansion of the strongest input without changing the second strongest input. (A) Histogram of the average rate of rise for all inputs (open) and the strongest inputs (closed). Line indicates the threshold for ‘strong inputs’. (B) For every cell the inputs are incrementally sorted by their average EPSP rate of rise. Diamonds indicate the inputs associated with a prespike. Colors indicate the ages as in C. Note the root scale of the ordinate. (C) Cumulative distributions of the competition index, i.e. the ratio of the rates of rise of the second strongest to the strongest input. (D) The rate of rise of the second strongest input against the strongest. Markers correspond to individual principal cells. Closed markers indicate that both inputs were identified based on spontaneous activity. Line shows identity line. Colors indicate age as in C.

### Strong inputs and prespikes

The effects of electrical stimulation of the calyx of Held synapse have been extensively studied in slices (Borst & Soria van Hoeve, 2012), but equivalent *in vivo* recordings have not yet been reported. In nine cells we could activate an input associated with an EPSP >10 V/s that could easily be distinguished from the other EPSPs. These inputs could be activated with a threshold current of 0.24 ± 0.09 mA (mean ± SD; n = 9 inputs in 9 cells, range 0.12 – 0.40 mA), although the reliability of activation with current above threshold varied between pups (83 ± 12% of the stimulations, mean ± SEM; range 59-100%). The average latency of the input’s EPSP was 2.1 ± 0.3 ms (mean ± SD, n = 9 inputs in 9 cells, range 1.6 – 2.5 ms) with little jitter between trials (SD/mean: 2.3 ± 0.9%, n = 9 inputs, range 1.0-3.8%). Given the distance of 500-1000 μm between the stimulation and recording electrode and a synaptic delay of ^~^0.5 ms, the propagation speed can be estimated to be in the order of 1 m/s. The AP travel time became less variable with development (n= 9 inputs in 9 cells, jitter vs. age, *r* = −0.75, −0.7 ± 0.2% per day, F_1,7_ = 9.0, *p* = 0.02). In addition, a decrease in latency (*r* = −0.7, −0.18 ± 0.07 ms/day, F_1,7_ = 5.6, *p* = 0.05) suggested a developmental increase in conduction speed. This increase is likely underestimated, as we positioned the stimulation electrode as contralaterally as was permitted by the cranial window to be able to separate the stimulus artefact from the responses. These developmental changes will contribute towards the precise timing of the calyx of Held in adult, hearing animals.

The calyx of Held can typically be identified in postsynaptic recordings by the presence of a prespike, a small deflection preceding the large EPSPs (Forsythe, 1994; Borst *et al*., 1995; Lorteije *et al*., 2009). Prespikes can already be detected in the first postnatal week (Crins *et al*., 2011; Sierksma *et al*., 2016). We therefore inspected every strong input whether a prespike was present (Figure 3A). In four of four cells >P5, six of ten cells of P4-5, and in none of six P2-3 cells large EPSPs were preceded by a prespike (Figure 3B). Hence, the presence of the prespike before a large EPSP was more likely at older ages (*r* = 0.6); a fit with a logistic function yielded a mid-point of 4.1 ± 0.4 days postnatally (mean ± SD) and a steepness of 0.7 ± 0.4 days, suggesting that within two days most strong inputs will become associated with a prespike. The prespike can be considered to be a sensitive indicator of the currents that flow across the release face. As a consequence, this developmental switch may be explained by an increase in these currents. The size of these currents depends on apposition area (Hoffpauir *et al*., 2006; Holcomb *et al*., 2013), time course of the calyceal AP, which accelerates considerably during early development (Taschenberger & von Gersdorff, 2000; Sierksma & Borst, 2017), isochronicity of the presynaptic currents, and the subcellular distribution of presynaptic ion channels. In addition, the coupling depends on synaptic cleft resistivity and postsynaptic admittance (Savtchenko, 2007). The rapid change in most of these properties during this period should underlie the emergence of the prespike.

**Figure 3.**
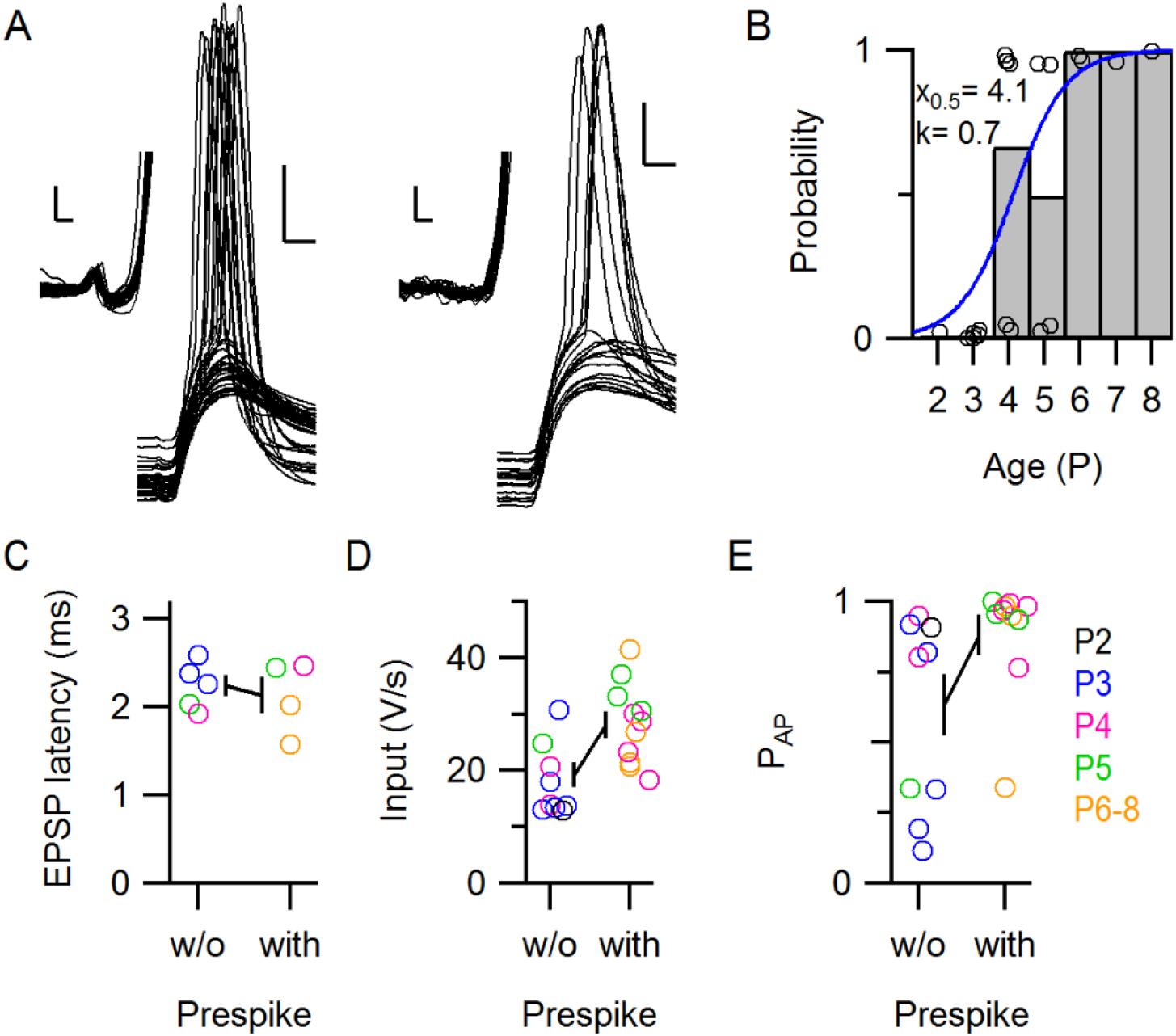
Comparison of strong inputs with and without prespikes. (A) Large EPSPs of a strong input with (left; P6) and of a strong input without (right; P5) a prespike. EPSPs were aligned on their onset. The insets show the same traces with their V_m_ offsets subtracted. Scale bars indicate 10 mV and 1 ms, and scale bars in the inset indicate 0.5 mV and 0.2 ms. (B) Developmental changes in the presence of a prespike preceding a large EPSP. Circles indicate cells where a large EPSP was recorded. Bars represent age average. Blue line represents the fit of the logistic function with the values reported in the graph. (C-E) Comparison of EPSP latency, average EPSP rate of rise and probability of eliciting postsynaptic APs (P_AP_) for strong inputs with or without a prespike. In C only evoked large EPSPs were analyzed. Circles indicate individual cells. Line represents the average and SEM.

As we observed strong inputs with and without prespike, we asked whether their properties were different. For the inputs that were electrically stimulated, the strong inputs with and without a prespike had a latency of 2.1 ± 0.4 ms and 2.2 ± 0.3 ms, respectively (Figure 3C, n = 4 and 5 inputs in 4 and 5 cells, respectively; *t*_7_ = 0.5, *r* = −0.2, *p* = 0.7). For the EPSP rate of rise, amplitude and the efficacy to trigger a postsynaptic AP (P_AP_), we included the strong inputs that were identified in the spontaneous activity. The EPSP rate of rise was 28 ± 2 V/s versus 19 ± 2 V/s (Figure 3D, age correction, n = 20 inputs in 20 cells, *F*_2,17_ = 5.3, effect of prespike, 5 ± 4 V/s, *t*_17_ = 1.2, *p* = 0.3) and had a coefficient of variation in the range of 7-38%. EPSP amplitude, which did not significantly correlate with age (*r* = 0.2), was 19.6 ± 1.1 mV versus 15.6 ± 1.5 mV (n = 10 and 10 inputs in 10 and 10 cells; *t*_18_ = 2.1, *r* = 0.45, *p* = 0.06). As these EPSPs typically triggered an AP (75 ± 31%, mean ± SD; range 11-100%), their amplitudes are underestimated. The fraction of EPSPs that triggered a postsynaptic AP was 0.88 ± 0.22 versus 0.63 ± 0.34 (Figure 3E, n = 20 inputs in 20 cells, rank-transform and age correction, *F*_2,17_ = 5.4, effect of prespike: 7 ± 3, *t*_17_ = 2.6, *p* = 0.02). Overall, prespike-associated inputs triggered postsynaptic activity more effectively. Still, some inputs that did not generate a detectable prespike were comparable to the calyceal inputs in strength and efficacy, as we observed before (Sierksma *et al*., 2016). Do these strong inputs without prespikes correspond to calyx of Held synapses? To answer this question, we looked at the synaptic structure of the recorded cells.

### The active zone protein Piccolo in the neonatal auditory brainstem

The dramatic developmental change in the glutamatergic innervation of MNTB neurons can be visualized by VGluT immunolabeling (Rodríguez-Contreras *et al*., 2008). However, VGluT does not mark release sites or postsynaptic densities. Wherease some postsynaptic density proteins appear delayed compared to VGluT labeling (Soria Van Hoeve & Borst, 2010), Piccolo, an active zone protein, was recently shown to be already present in P9 calyces (Dondzillo *et al*., 2010). We tested whether Piccolo could be used to visualize the release face of terminals within the first neonatal week. Already at P3-4, Piccolo-labeling was found throughout the superior olivary complex (SOC) in most auditory nuclei (*n* = 8 rats; Figure 4A). Although the staining was not particularly strong in the MNTB, it clearly co-labeled large perisomatic structures containing VGluT (Figure 4B). As early as P2, the Piccolo labeling could delineate an edge of the larger VGluT-clusters (Figure 4C, *n* = 2 of 5 P2 animals). Piccolo puncta were also found outside the somatic regions of the MNTB, and a Piccolo punctum was occasionally found within the soma of a principal cell (Figure 4C). Generally, when imaged with confocal microscopy, the Piccolo edges appeared to be composed of individual spots, which were close to the diffraction limit (Figure 4C-D). We turned to structured-illumination microscopy (Gustafsson, 2005), which revealed that the Piccolo clusters were indeed composed of multiple spots (Figure 4D). These spots were also found in biocytin-filled calyces, indicating that Piccolo can be present in the active zones of the calyx of Held at P5 (*n* = 11 of 12 calyces; Figure 4E). Piccolo thus identifies the release face of synapses and thereby facilitates the identification of the target neuron in the case of axosomatic VGluT clusters (Figure 4C-D). However, dendritic synapses could not be reliably identified without labeling the postsynaptic neuron.

**Figure 4.**
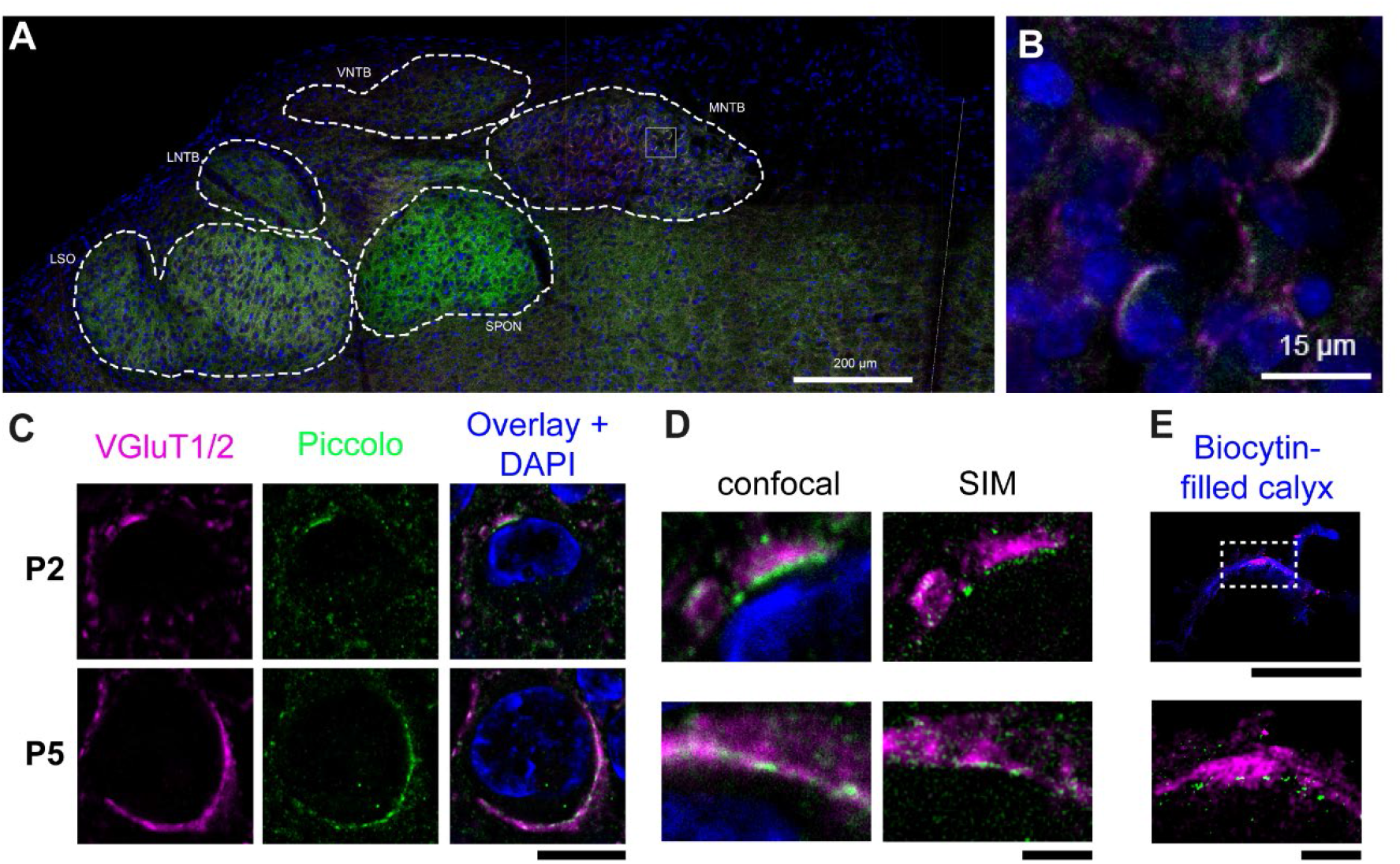
Developmental expression of Piccolo in the MNTB. (A) Fluorescent labeling of Piccolo (green), VGluT (magenta) and DAPI (blue) in the SOC of a P4 rat. Midline is indicated with a white line. Auditory nuclei are delineated with dashed lines. (B) Higher magnification of the grey box in A showing principal cells of the MNTB with large perisomatic clusters of VGluT and Piccolo. (C) A P2 and P5 principal cell with fluorescent labeling of VGluT, Piccolo, and VGluT + Piccolo + DAPI with confocal microscopy. (D) Comparison of confocal microscopy and SIM of perisomatic clusters of Piccolo and VGluT of the P2 principal cell shown in C (top), and of the P5 principal cell shown in C (bottom). (E) SIM image of a P5 biocytin-filled calyx (top, blue) with VGluT and Piccolo, revealing active zone-like structures in the Piccolo labeling (bottom). Non-calyceal labeling was masked. Scale bars indicate 200 μm in A, 15 μm in B, 10 μm in C, 2 μm in D, and 10 and 1 μm in E. Images in C and D were background-subtracted. Abbreviations: LSO, lateral superior olive; MNTB, medial nucleus of the trapezoid body; P, postnatal; SIM, structured-illumination microscopy; SOC, superior olivary complex; SPON, superior periolivary nucleus; VGluT, vesicular glutamate transporter 1 and 2; VNTB, ventral nucleus of the trapezoid body.

### Structure-function correlation within and across cells

From the 32 recorded cells, we recovered 20 cells with immunolabeling for VGluT (1 P2, 7 P3, 8 P4, 2 P5, 1 P6, 1 P7). For these cells we segmented the soma and the VGluT clusters surrounding the soma using a new, semi-automatic procedure (see Methods, Figure 5A). This segmentation takes into account that a VGluT cluster may have multiple contact points with the soma by clustering connected voxels. A VGluT cluster is thus defined as a contiguous structure of VGluT-positive voxels and VGluT clusters, by definition, were not in contact with each other. The total somatic surface was 620 ± 230 μm^2^ (mean ± SD, n = 20), which is comparable to earlier estimates (Sommer *et al*., 1993; Hoffpauir *et al*., 2006). We observed that multiple VGluT clusters contacted the postsynaptic cell, of which most had a postsynaptic contact area <3 μm^2^ (Figure 5B-C). The total postsynaptic area that was covered by VGluT ranged from 2.4 μm^2^ to 76.5 μm^2^. As calyces can cover >50% of the soma (Sätzler *et al*., 2002; Hoffpauir *et al*., 2006), our segmentation of VGluT thus may give us only a lower-bound approximation of its size. For each cell we identified the largest and second largest VGluT cluster. The contact area of the largest cluster on the postsynaptic cell ranged from 0.4 μm^2^ to 59.7 μm^2^ (13 ± 16 μm^2^, mean ± SD). For the second largest cluster it ranged from 0.3 μm^2^ to 10.0 μm^2^ (2.4 ± 2.8 μm^2^, mean ± SD). For 3 cells the Piccolo labeling failed. For 9 cells we could see clear Piccolo labeling coinciding with the largest VGluT cluster directed towards the biocytin-filled cell. For 8 cells VGluT clusters surrounding the biocytin-labeled soma did not show clear Piccolo labeling, but Piccolo coincided with VGluT in the surrounding tissue. The signal-to-noise ratio of the reconstruction did not allow a more in-depth analysis of the Piccolo clusters.

**Figure 5.**
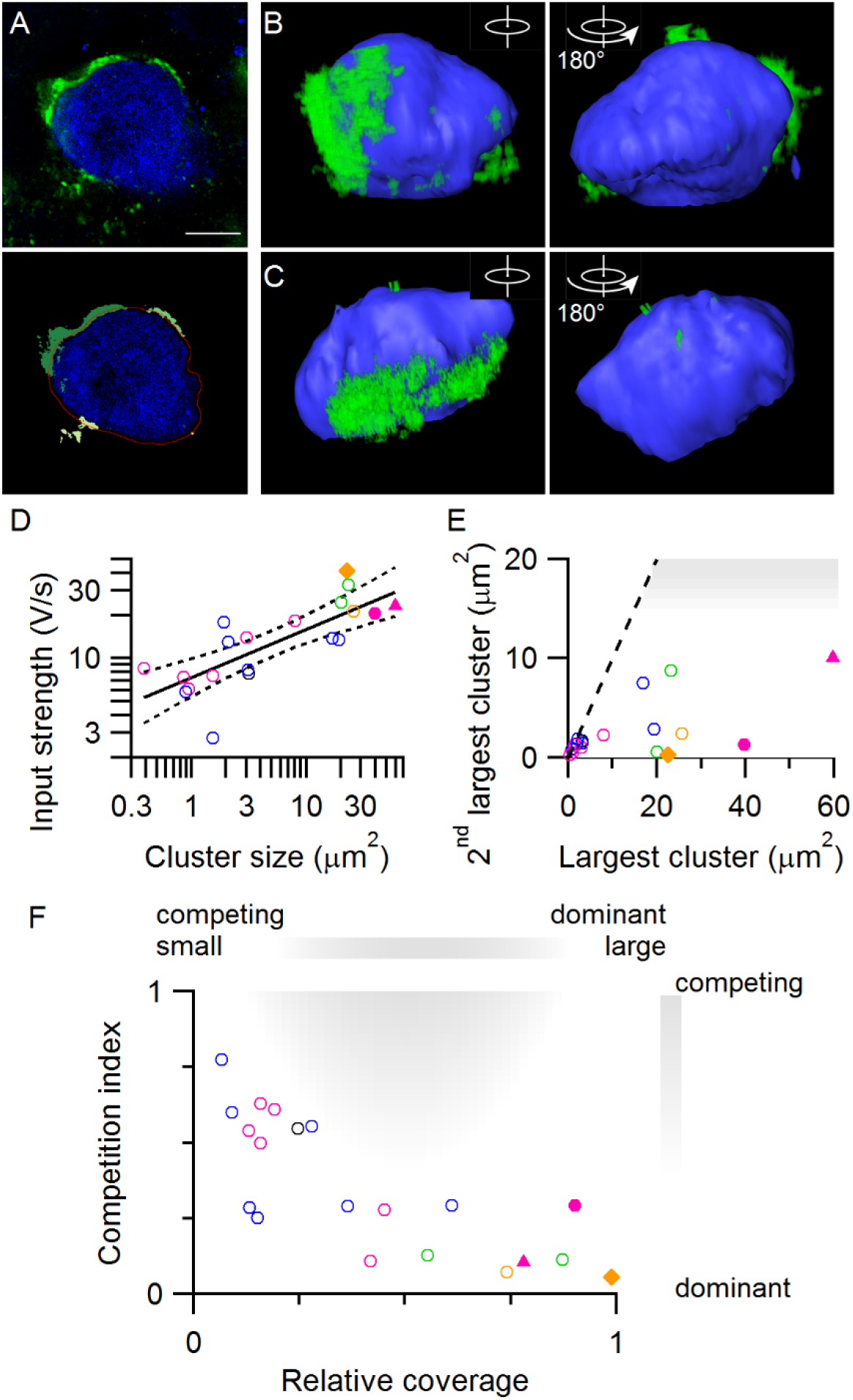
Contact area of perisomatic VGluT clusters correlates with synaptic strength *in vivo*. (A) Example of segmentation of VGluT clusters around the soma. The image before (top) and after segmentation (below). Blue: biocytin; Green: VGluT labeling. In the bottom image different VGluT clusters are depicted by different shades of green. Red line indicates contour of the cell after dilation. For this cell 1 input was associated with a prespike. (B-C) 3D reconstructions of somata (blue) with VGluT clusters (green). Dendrites were manually removed. Right image is the same soma after 180 degrees rotation. Postsynaptic cell in B did not have a prespike-associated input. The VGluT cluster in C was manually separated from a nearby cluster that terminated on another cell. (D) Relation between input strength of strongest input and contact area of the largest VGluT cluster for each cell. Line indicates linear fit of log-transformed variables. Dotted line indicates 95% confidence intervals. (E) Relation between area of largest and second largest VGluT cluster. Shaded area: possibly multi-calyceal innervation. (F) Competition index against relative coverage. Competition index equals the second strongest input relative to the strongest input. Relative coverage equals the contact area of the largest VGluT cluster relative to the total area contacted by VGluT clusters. Shaded area indicates region where data points with multi-calyceal competition are expected to appear. Colors in D-G indicate age as in previous figures. Triangle indicates the data point that corresponds to the cell shown in A. Closed circle and diamond indicate data points corresponding to the cells shown in B and C, respectively.

If the size of the VGluT cluster is a useful estimate of the size of the terminal, then it should predict the presence and amplitude of a prespike in our *in vivo* recordings. The size of the VGluT cluster was moderately correlated with the presence of a prespike (ρ = 0.63). The largest VGluT cluster was 28 ± 19 μm^2^ (mean ± SD) for the 5 cells in which we recorded a prespike. This was significantly larger than in the 15 cells without a prespike (8 ± 11 μm^2^, mean ± SD, *t*_18_ = 3.3, *p* = 0.004). Next, we compared the prespike peak-to-peak amplitude with the size of the largest VGluT cluster. We excluded one of the recordings, in which the prespikes had an amplitude of 15 mV. In this recording a small patch of calyceal membrane may have been present within the electrode tip (Forsythe, 1994). As the calyx was not stained for biocytin, the membrane of the calyx was apparently not damaged by the pipette. For the remaining prespikes the average amplitudes were 0.11, 0.26, 0.30 and 0.37 mV, which corresponded with values of 19.2, 23.1, 25.6 and 59.7 μm^2^ for the largest VGluT cluster (*r* = 0.8). Because of the small number of observations, this correlation did not reach significance (F_1,2_ = 3, *p* = 0.2). Also, we observed VGluT clusters with similar contact area among the cells from which we did not record a prespike. Altogether, the size of the largest VGluT cluster may indicate the presence of a large terminal, but it is unlikely to reveal the entire contact area made by the terminal.

As synaptic strength depends more on glutamate release than on terminal size, the observed VGluT cluster may still be a good approximation of synaptic strength. We therefore compared synaptic strength with the contact area of VGluT clusters (Figure 5D). The size of the strongest input was correlated to the size of the largest VGluT cluster (log-transform, *r* = 0.8, F_1,18_ = 26, *p* < 0.001). The structure-strength relation could be approximated by EPSP (V/s) = β area^α^ with β = 6.8 ± 0.9 V s^−1^ μm^−2α^ and α = 0.35 ± 0.06. Thus, the input strength was correlated with the cube root of VGluT area. For the older pups (2 P5, 1 P6 and 1 P7 cell) the structure-function relation was underestimated; three of four synapses were stronger than expected from their apposition area. This may indicate that there is a developmental change in the structure-function relation or that the total vesicle-loaded contact of giant synapses is underestimated as the largest VGluT cluster may be only a small part of the synapse.

Calyces that innervate a multi-calyceal principal cell are filled with synaptic vesicles (Holcomb *et al*., 2013), and would thus be expected to be observed with VGluT labeling. The different calyces were observed to each occupy a contiguous area, sometimes at opposite sides of the target neuron (Holcomb *et al*., 2013). This type of multi-calyceal innervation was not observed here (Figure 5B, C and E). As we cannot be sure whether individual VGluT clusters are independent inputs or belong to the same synapse, we cannot use the second largest VGluT clusters to relate to the physiology. Indeed, the size of the second largest VGluT cluster was not correlated with the size of the second strongest input (*r* = −0.1, F_1,16_ = 0.3, *p* = 0.6). Nonetheless, for multi-calyceal innervation we would have expected to observe two large VGluT clusters of comparable contact area (Figure 5E). No pair of VGluT clusters was unambiguously identified as such.

As we cannot conclusively assign a VGluT cluster to individual inputs, it may still be that the second calyx contacted the cell via multiple clusters. We therefore quantified how the largest VGluT cluster relates to all the other clusters that contact the soma of the same target neuron as the contact ratio of the contact area of the largest VGluT cluster to the total somatic surface contacted by VGluT. When the contact ratio approaches 100% the largest VGluT cluster is the only cluster that contacts the soma of the target neuron, and values <100% indicate the presence of additional VGluT clusters which may or may not belong to the same terminal. The contact ratio ranged from 9-98% (41 ± 30%, mean ± SD) and was negatively correlated to the competition index (Figure 5F, F_2,17_ = 10, *p* = 0.0012, age correction, ρ = −0.7). With these two measures, the competition index and the contact ratio, we could have identified multi-calyceal competition as (*i*) two terminals that cover about an equal size of the soma, *i.e*. a contact ratio of ^~^0.5, and (*ii*) two inputs of similar strength, giving a competition index close to 1 (shaded area in Figure 5F). Such combinations were not observed. We thus conclude that we did not find strong evidence in support of multi-calyceal competition. In addition, there were no large discrepancies between the structure and function of the developing calyx of Held synapse. This suggests that in the developing rat MNTB a large VGluT cluster likely indicates the presence of a strong input that is clearly stronger than the other inputs of its target neuron.

## Discussion

In this study we combined *in vivo* electrophysiology with immunolabeling of the developing calyx of Held synapse. With afferent stimulation we identified multiple inputs for each principal neuron. Increasing stimulation levels showed either a graded increase or a sudden, large jump in the rate of rise of the response. The strongest inputs across cells showed a shorter latency. The strongest inputs were typically at least twice as strong as the other inputs. This divergence increased during development with little change in the strength of the second strongest input. We found that in the first postnatal week Piccolo is already present within perisomatic VGluT clusters, which are putative calyceal terminals. The contact surface of the largest VGluT cluster correlated with the strength of the strongest input *in vivo*. Furthermore, the relative coverage of the largest VGluT cluster correlated with the relative strength of the strongest to the second strongest input. Overall, VGluT-positive structures correlated with synaptic strength *in vivo*. The association of a strong input with a prespike became more likely during development. Prespike-associated inputs were more effective in triggering postsynaptic activity, but some inputs without a prespike had similar properties.

### Calyceal competition during development?

The prevalence of multiple calyces on a principal MNTB neuron is a subject of debate. On the one hand, multiple strong inputs have been observed in slice recordings (Bergsman *et al*., 2004; Hoffpauir *et al*., 2006; Hoffpauir *et al*., 2010), and structural evidence for the presence of multiple large terminals on rodent MNTB neurons has also been obtained (Wimmer *et al*., 2004; Hoffpauir *et al*., 2006; Holcomb *et al*., 2013; Matho, 2013; Milinkeviciute *et al*., 2019). Although only a minority (up to 10%) of neurons is persistently innervated by multiple calyces (Holcomb *et al*., 2013; Matho, 2013; Milinkeviciute *et al*., 2019), as many as half of the principal neurons are putatively contacted by multiple large terminals during development (Holcomb *et al*., 2013). On the other hand, to our knowledge not a single conclusive example of multi-calyceal innervation has been reported for *in vivo* studies of the adult MNTB. Multi-calyceal innervation could have been observed as inputs associated with prespikes that do not obey the (relative) refractory period. Neither our *in vivo* recordings nor our VGluT labeling revealed multi-calyceal innervation in this study. Multi-calyceal competition would be expected to involve an input strength that is within a similar range. This was observed in our earlier study where three out of 132 cells had two strong inputs that could be separated based on the presence or absence of a prespike and that defied the refractory period (Sierksma *et al*., 2016). However, we cannot exclude that one of these inputs was dendritic. Even if we grant that these three cells demonstrate multi-calyceal competition, we identify, from our earlier (3/132 cells) and the current (0/32 cells) experiments, <10% of rat principal neurons with multi-calyceal innervation during development. These values are comparable to the 1% reported in a pre-hearing mouse study (Kronander *et al*., 2019) and to our earlier study in P9 rats, in which 1 of 86 cells showed two large, perisomatic VGluT clusters for (Rodríguez-Contreras *et al*., 2006). Even though we cannot exclude having missed calyces due to low VGluT expression, we conclude that multi-calyceal competition is unlikely to significantly contribute to the development of the rat calyx of Held synapse.

The presence of competition has been evaluated by the (relative) contact surface (Holcomb *et al*., 2013) or by the (relative) strength of the inputs (Hoffpauir *et al*., 2010; Sierksma *et al*., 2016). Here, we compared a structural and a functional measure for competition, and generally observed a good match between the two. Although immunolabeling did not allow the unambiguous identification of different inputs based on the VGluT clusters, we did observe a clear match between VGluT relative coverage and the competition index, indicating a link between the physiological and morphological development of the MNTB synapses. In addition, it may point to a possible period of competition prior to calyx formation as we only observed inputs with competing strength at P2-3, when the relative coverage of the largest VGluT cluster was still low and only few calyces have already formed (Morest, 1968; Kandler & Friauf, 1993; Rodríguez-Contreras *et al*., 2008).

For two other giant, mono-innervating synapses, the neuromuscular junction (NMJ) and the cerebellar climbing fiber-Purkinje cell synapse (CF-PC), different types of competition have been described. In the case of NMJ, before mono-innervation is established, one input becomes >4 fold stronger than the other ones by both an increase of the strongest and a decrease in the strength of the other inputs (Colman *et al*., 1997). As NMJ inputs compete for a small patch of the muscle membrane (Sanes & Lichtman, 1999; Darabid *et al*., 2014), the expansion of one NMJ necessarily involves the shrinkage of another, as reflected in the opposing changes in strength. In contrast, CFs compete at the soma for access to the apical dendrite (Watanabe & Kano, 2011). A fourfold strengthening of the strongest input was observed during development (Hashimoto & Kano, 2003) while other inputs remained constant (Bosman *et al*., 2008). Subsequently, one of the immature CFs is allowed to translocate to the apical dendrite (Hashimoto *et al*., 2009; Watanabe & Kano, 2011), ending the competition (Carrillo *et al*., 2013). Our observations seem to reflect a CF-PC-like competition. We observed that the strongest input became >4 times stronger than the other inputs between P3-5 without a substantial change for the second strongest. This may indicate that they do not compete for overlapping synaptic domains (Holcomb *et al*., 2013). In addition, the strongest input already reached this relative strength while its coverage still seemed to be low, suggesting that the increase in strength precedes and instructs the expansion (Hoffpauir *et al*., 2010). In contrast, the combination of a large VGluT cluster with a high competition index was not observed, suggesting that somatic expansion and/or VGluT filling of the terminal (Rodríguez-Contreras *et al*., 2008) represents a relatively late maturation stage.

An important role for exuberant connectivity seems to be to provide sufficient candidate precursors to ensure that each principal neuron will eventually end up being innervated by a calyx of Held from a globular bushy cell that innervates only a limited number of other principal neurons. As the molecular mechanisms underlying this process are gradually being elucidated (Hsieh *et al*., 2010; Nakamura & Cramer, 2011; Ehmann *et al*., 2013; Michalski *et al*., 2013; Xiao *et al*., 2013; Körber *et al*., 2014; Yu & Goodrich, 2014; Nothwang *et al*., 2015; Willaredt *et al*., 2015; Dimitrov *et al*., 2016), to what extent postsynaptic activity plays a role in this selection process is still unclear. Without concurrent activity of other inputs, non-calyceal inputs are unable to evoke postsynaptic APs, as they elicit postsynaptic firing almost exclusively by EPSP summation (Sierksma *et al*., 2016 and this study). In the fruit fly postsynaptic activity stimulates the release of bone morphogenetic protein (BMP) homologue which promotes active zone development and NMJ growth (Berke *et al*., 2013; Berke *et al*., 2020). Mice conditionally deleted for BMP receptor 1 and 2 show deficits in transmitter release and smaller calyces (Xiao *et al*., 2013) in addition to a persistence of axonal branches with calyces and increased multi-calyceal innervation (Kronander *et al*., 2019). Similarly, calyces fail to form when synaptic transmission is impaired by genetic deletion of dynamins (Fan *et al*., 2016). These phenotypes may reflect a need for postsynaptic activity in calyx development.

Following calyx formation, most non-calyceal inputs are gradually pruned over a prolonged period (Morest, 1968; Kandler & Friauf, 1993; Rodríguez-Contreras *et al*., 2008), whereas some may persist (Guinan & Li, 1990; Hamann *et al*., 2003). This pruning phase may be mediated by microglia (Milinkeviciute *et al*., 2019).

### Structure-function relation for the developing calyx of Held synapse

The presence of a prespike indicates the innervation by a giant terminal (Forsythe, 1994). In this study four cells with a prespike-associated input had a large VGluT cluster and one cell did not. Possibly in the latter case the terminal was larger than revealed by VGluT labeling (Rodríguez-Contreras *et al*., 2008). Conversely, four cells were observed where the presence of a large VGluT cluster did not match with a prespike-associated input. In these cases we may have missed small prespikes due to the capacitive filtering by the soma, and the prespikes may have been detectable in voltage-clamp recordings. There are also a number of other biological factors that influence the prespike amplitude (see Results), but experimental observations on the relation between presynaptic AP and the prespike have so far been qualitative (Forsythe, 1994; Borst *et al*., 1995; Wang *et al*., 2017). Whereas the presence of a clear prespike seems to be good evidence for the presence of a giant somatic terminal, experimental and computational studies on the prespike (Savtchenko, 2007) based on exact morphology (Spirou *et al*., 2008) are needed to allow a more precise estimate for the minimal conditions under which a calyx generates a prespike.

From our structure-function analyses we found that VGluT labeling closely follows the strength of the strongest input. This VGluT cluster represents the total pool of glutamatergic vesicles in the calyx of Held (Rodríguez-Contreras *et al*., 2008). This pool likely covers multiple active zones (Sätzler *et al*., 2002; Taschenberger *et al*., 2002; Hoffpauir *et al*., 2006; Dondzillo *et al*., 2010), which we observed with the Piccolo labeling. The relation between the size of this pool and synaptic strength is complex, depending on among others quantal size, release probability, and number of release sites (Atwood & Karunanithi, 2002; Schneggenburger & Forsythe, 2006; Borst & Soria van Hoeve, 2012). We found a cube root relation between VGluT contact surface and EPSP rate of rise, suggesting that the increase in vesicle pool is relatively ineffective in increasing synaptic strength. An important factor may have been that EPSC-EPSP relation saturates for large EPSCs (Lorteije *et al*., 2009; Sierksma *et al*., 2016), which would lead to a sublinear relation between quantal content and EPSP size. During early development quantal size is stable (Chuhma & Ohmori, 1998; Taschenberger & von Gersdorff, 2000) or may slightly decrease (Rusu & Borst, 2011), while the active zones and postsynaptic densities become more numerous and segregate into smaller clusters (Taschenberger *et al*., 2002; Hoffpauir *et al*., 2006; Soria Van Hoeve & Borst, 2010). Other possible factors include changes in release probability, homeostatic plasticity of postsynaptic excitability (Hoffpauir *et al*., 2010; Rusu & Borst, 2011; Sierksma *et al*., 2016) or a contribution of dendritic inputs (Rodríguez-Contreras *et al*., 2008; Holcomb *et al*., 2013).

For the study of the relationship between structure and function, the calyx of Held synapse has the advantage of an axosomatic, (putative) one-to-one innervation facilitating the identification of the strongest input *in vivo* with the largest cluster *ex vivo*. An even more direct relationship might be possible following simultaneous electrophysiology and calcium imaging *in vivo* (Winnubst *et al*., 2015). Subsequent reconstruction of the synapses after *in vivo* imaging (Liang *et al*., 2018) would allow a structure-function description for multiple synapses of a single cell. Such studies may further define the relationship between morphological and functional properties of the precursors of the calyx of Held. Finally, longitudinal imaging might reveal some of the dynamics associated with the formation of the calyx of Held and the interactions with surrounding axons (Rodríguez-Contreras *et al*., 2008).

## Additional information

### Contributions

JGGB and MCS designed experiments. MCS performed *in vivo* electrophysiology and imaging experiments, analyzed the electrophysiology and the images. JAS performed imaging experiments and imaging analysis. ABH, JAS and JGGB designed imaging analysis. MCS drafted the manuscript. JAS, ABH and JGGB revised it critically for important intellectual content. All authors approved the final version of the manuscript and agree to be accountable for all aspects of the work in ensuring that questions related to the accuracy or integrity of any part of the work are appropriately investigated and resolved. All persons designated as authors qualify for authorship, and all those who qualify for authorship are listed.

## Acknowledgements

We are very grateful for the help of Elize Haasdijk and Celina Glimmerveen, who performed immunolabeling. We thank Marcel van der Heijden, Peter Bremen and Aaron Wong for advice on the analyses.

